# Discovery of Epigenetically Silenced Tumour Suppressor Genes in Aggressive Breast Cancer Through a Computational Approach

**DOI:** 10.1101/2025.01.31.635071

**Authors:** Anne-Laure Vitte, Florent Chuffart, Emmanuelle Jacquet, Eleni Nika, Mireille Mousseau, Ina Jung, Séverine Tabone-Eglinger, Thomas Bachelot, Sophie Rousseaux, Saadi Khochbin, Ekaterina Bourova-Flin

## Abstract

Breast cancer is characterized by genetic and epigenetic deregulations, leading to aberrant expression of tissue-specific genes that are normally silent in healthy breast tissue. Our previous work identified the embryonic stem cell-specific gene *DNMT3B*, a *de novo* DNA methyltransferase, as aberrantly activated in breast cancer, correlating with aggressive tumour behaviour and high relapse risk, regardless of molecular subtype. Through integrative bioinformatic analyses of DNA methylation and transcriptomic data, we identified 154 genes downregulated via *DNMT3B*-driven promoter hypermethylation, many of which are associated with high relapse risk. Notably, the tumour suppressor gene *GATA3* emerged as a primary target of functional inactivation through either loss-of-function mutations or *DNMT3B*-controlled hypermethylation, in a mutually exclusive manner.

Both mechanisms of *GATA3* inactivation were associated with similar molecular signatures linked to tumour progression, increased malignancy, and poorer prognosis. However, distinct differences were observed, with immune- and inflammation-related genes enriched in *GATA3* hypermethylation cases but depleted in mutation-driven silencing. Additionally, our analysis uncovered other potential tumour suppressor genes epigenetically repressed in aggressive breast cancers. These findings underscore a broader role of *GATA3* inactivation beyond genetic alterations and suggest therapeutic opportunities to target epigenetically silenced tumour suppressors in aggressive breast tumours.

## Introduction

Breast cancer is the most common cancer in women, with 2.3 million new cases and 685 000 deaths worldwide in 2020. It is currently stratified into subtypes on the basis of the expression of biomarkers, notably the receptors for oestrogen and progesterone (ER, PR), human epidermal growth factor receptor2 (HER2), and Ki-67 (Sørlie et al., 2003; Cancer Genome Atlas Network, 2012; Parker et al., 2023). Luminal A tumours (40-50% of cases) express ER and PR, are HER2-negative and have low levels of Ki-67. This tumour subtype is less aggressive, responds well to hormone therapy and has a five-year survival rate of over 90%. Luminal subtype B (15-20%) expresses less ER and PR, may be HER2-positive and has high levels of Ki-67. More aggressive, these tumour types require hormone therapy, chemotherapy and HER2-targeted therapies, with a survival rate of around 80-85%. HER2-enriched breast cancers (10-15%) are ER/PR negative, and very aggressive, but responds well to HER2-targeted therapy (trastuzumab) and chemotherapy, with a survival rate of 80-85%. Triple-negative breast cancer (TNBC) (15-20%) lacks ER, PR and HER2 (Smolarz et al., 2022). Very aggressive with limited treatment options, TNBC has a survival rate of 60-70% (Fayaz et al., 2019; Baranova et al., 2022). Immunophenotyping enables personalised treatment based on tumour characteristics. In general, therapies targeting HER2 and hormonal pathways enhance treatment outcomes. Additionally, emerging treatments targeting biomarkers such as PD-L1 (for TNBC) and the androgen receptor offer promising new therapeutic options (Wang and Minden, 2022; Papalexis et al., 2024).

In breast cancer, as in all cancers, genetic and epigenetic deregulations can result in out-of-context expressions of a set of normally silent tissue-specific genes, including the genes predominantly expressed in testis, placenta and embryonic stem cells. An aberrant activation of tissue-specific genes can empower tumour cells with new properties, related to enhanced proliferation and metastatic spread, and leading to a poor survival prognosis. For this reason, off-context activations represent a promising source of novel prognostic biomarkers and therapeutic targets in breast cancer. Following an original approach based on the identification of off-context tissue-specific gene activation in breast cancer, we recently identified the *de novo* DNA methyltransferase *DNMT3B*, along with four other genes, as markers of highly aggressive breast cancer subtypes (Jacquet et al., 2023).

The present study confirms that expression of the *DNMT3B* gene alone, independently of the other marker genes, is also associated with a poor prognosis in breast cancer patients. The molecular mechanisms associated with *DNMT3B* expression were then further investigated by comparing DNA methylation levels in high-*DNMT3B*-expressing tumours (DNMT3B+) versus low-*DNMT3B*-expressing tumours (DNMT3B-) samples in the TCGA-BRCA public dataset, which combines expression and methylome data from the same breast cancer tumours, and then confirmed in an independent cohort. Identification of differentially methylated regions (DMR) and of the genes associated with these regions, led to the identification of a subset of specific genes, including *GATA3*, targeted by *DNMT3B*-dependent epigenetic repression. Interestingly, further investigations showed that *GATA3* silencing, either caused by aberrant hypermethylation or by DNA alterations, is also associated with a poor prognosis. Gene Set Enrichment Analysis (GSEA) of both situations relative to control breast cancers expressing *GATA3*, highlighted shared associated gene expression signatures, including cancer aggressiveness and cell proliferation, confirming that *GATA3* is indeed a powerful tumour suppressor gene. However, these analyses also showed that the genes involved in inflammatory and immune responses, which are highly enriched in the group of methylation-dependent *GATA3* silencing, are significantly depleted in the group of patients where *GATA3* is silenced through genetic alterations. Finally, this approach also highlighted a number of potential tumour suppressor genes that are inactivated by ectopic *DNMT3B* expression, and thus provided a novel insight into epigenetically-driven aggressive breast cancers.

## Results

### Out-of-context activation of the embryonic stem cell/germ cell-specific gene, *DNMT3B*, is significantly associated with a poor prognosis in breast cancer

As *DNMT3B* is a *de novo* methyltransferase, we hypothesised that its contribution to the increased malignancy of the tumours might be due to its ability to silence some critical tumour suppressor genes following an ectopic DNA-methylation-dependent silencing of these genes. Using the RNA-seq data available in the GTEX and NCBI datasets, we calculated the expression profile of the gene *DNMT3B* in non-tumour tissues. As shown in Figure 1A, *DNMT3B* is highly expressed in embryonic stem cells and has a moderate expression level in testis. In other somatic tissues, this gene is expressed at low levels or not at all. In particular, in healthy breast tissue, its expression levels correspond to the background signal. However, *DNMT3B* can be aberrantly activated in breast cancer. Figure 1B-C show the expression levels of *DNMT3B* in non-tumour and in tumour samples of the TCGA-BRCA dataset, in the total cohort, and in different molecular subtypes: luminal-A, luminal-B, HER2-enriched and basal-like. *DNMT3B* is significantly over-expressed in tumour samples compared to non-tumour breast tissues. Its level of expression is higher in HER2-enriched and basal-like molecular subtypes, which are generally considered to have a poor prognosis, than in luminal-A and luminal-B molecular subtypes of breast cancer, which have a better chance of survival. Indeed, survival analysis performed between two groups of low (DNMT3B-) and high (DNMT3B+) expression in eight independent breast cancer datasets revealed a significant and reproducible association of the expression level of this gene with the probability of disease-free survival, consistently predicting a higher risk of relapse in the DNMT3B+ group compared to DNMT3B-(Figure 1D). In each cohort, the DNMT3B- and DNMT3B+ groups were defined using a threshold corresponding to the median expression level of *DNMT3B* in all tumour samples. The association with the probability of disease-free survival found to be statistically significant using the log-rank test in seven of the eight breast cancer datasets analysed. In the last dataset, GSE42568, the p-value obtained was not formally significant (p=0.293), probably due to the smaller sample size in this cohort, which was the smallest of all the cohorts analysed.

**Figure 1.**
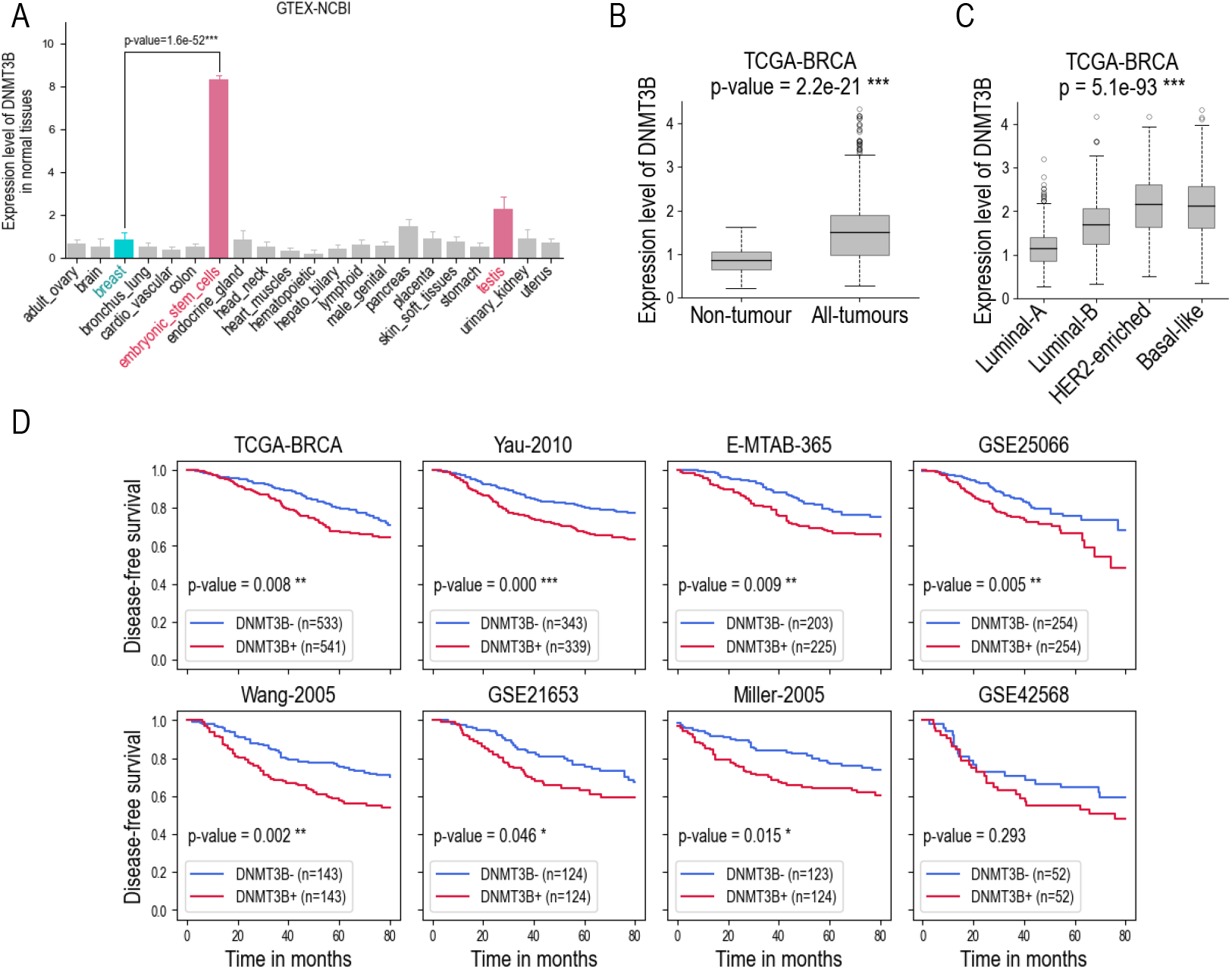
Aberrant activation of the embryonic stem cell-specific gene *DNMT3B* in aggressive forms of breast cancer. **A**: Expression levels of the gene *DNMT3B* in different healthy tissues in the GTEX-NCBI dataset. *DNMT3B* is predominantly expressed in embryonic stem cells, moderately expressed in testis and weakly expressed in other tissues. The difference in expression levels between embryonic stem cells and normal breast tissue is statistically significant (two-tailed t-test, p-value < 0.05). **B**: Expression levels of *DNMT3B* in non-tumour breast samples (labelled as “non-tumour” in the figure) versus tumour breast samples (labelled as “all tumours” in the figure) from the TCGA-BRCA dataset. The p-value corresponds to the two-tailed t-test comparing these groups. **C**: Expression levels of the gene *DNMT3B* in different molecular subtypes of breast cancer in the TCGA-BRCA dataset: luminal A, luminal B, HER2-enriched, and basal-like. The p-value corresponds to an ANOVA test across these subtypes. **D**: Kaplan-Meier survival curves across eight public datasets, calculated for two groups of patients based on *DNMT3B* expression levels: tumours with low *DNMT3B* expression (DNMT3B-, blue line) and tumours with high *DNMT3B* expression (DNMT3B+, red line). The threshold separating the DNMT3B- and DNMT3B+ groups correspond to the median expression level. P-values are calculated using the log-rank test and indicated by the following significance symbols: * for p < 0.05, ** for p < 0.01, and *** for p < 0.001.

### Out-of-context activation of *DNMT3B* is associated with a hypermethylation in CpG-rich promoter regions of several onco-suppressor genes

The gene *DNMT3B* encodes a DNA methyltransferase which regulates *de novo* DNA methylation, alongside with another enzyme *DNMT3A* (Bird, 1999; Okano et al., 1999). For instance, *DNMT3A* and *DNMT3B* enzymes can produce epigenetic modifications through DNA methylation of promoter regions of a specific set of genes. In principle, strong methylation of CpG islands in upstream regulatory regions of genes can be associated with the silencing of the corresponding genes. We hypothesised that overexpression of *DNMT3B* in breast cancer could lead to aberrant (ectopic) DNA methylation of multiple regions of the genome in tumour cells, which would abnormally reduce the expression of certain genes that are normally expressed in healthy breast tissue. With this aim in mind, we predicted that, in DNMT3B+ tumours, the hypermethylation associated with high levels of *DNMT3B* expression would not be randomly distributed across the genome, but could target specific regions containing critical genes, giving these tumours particular aggressive characteristics. To address this point, we carried out a joint analysis of the methylation and gene expression data available in the TCGA-BRCA dataset.

We used the methylation data of the TCGA-BRCA dataset containing 766 samples with annotations of 393 872 methylation sites provided by the Infinium Methylation450k array (Illumina, San Diego, CA, USA). We performed epigenome-wide association study (EWAS) to identify the differentially methylated probes between the groups DNMT3B+ versus DNMT3B-. The obtained p-values from the EWAS analysis were further processed with the “comb-p” software (Pedersen et al., 2012). The “comb-p” method allowed us to identify the differently methylated regions (DMR) forming series of adjacent probes associated with low p-values. Using the annotations of the Infinium Methylation450k array, these regions were characterised by their functional role (e.g., promoter region from 1Kb to 5Kb around the transcription start site (TSS), 5’ untranslated regions, exon, intron) and assigned to the corresponding genes. We found that the promoter regions 1Kb-TSS-5Kb or the 5’ untranslated regions (5’UTR) were significantly hypermethylated for 1939 genes with p-values obtained from the “comb-p” analysis lower than 1e-05. The list of these genes is given in Supp. Table S1. To investigate whether the DNA hypermethylation corresponding to the selected 1939 genes causes a down-regulation of their expression levels, we performed a differential expression analysis of the transcriptome between the DNMT3B+ and DNMT3B-groups in the TCGA-BRCA dataset. The volcano plot and heatmap in Figure 2A illustrate the expression of the genes respectively down- and up-regulated in DNMT3B+ versus DNMT3B-breast cancer samples with an absolute fold change of expression values above 1.5 and an adjusted Mann-Whitney p-value < 0.05. 646 and 606 genes were found to be downregulated and upregulated in DNMT3B+ versus DNMT3B-, respectively. At the intersection of the 1939 hypermethylated genes and the 646 genes downregulated in DNMT3B+, we identified 154 genes for which promoter or 5’UTR hypermethylation was systematically associated with down-regulation of their expression level (Figure 2B-E). The list of the 154 genes is detailed in Supp. Table S2.

**Figure 2.**
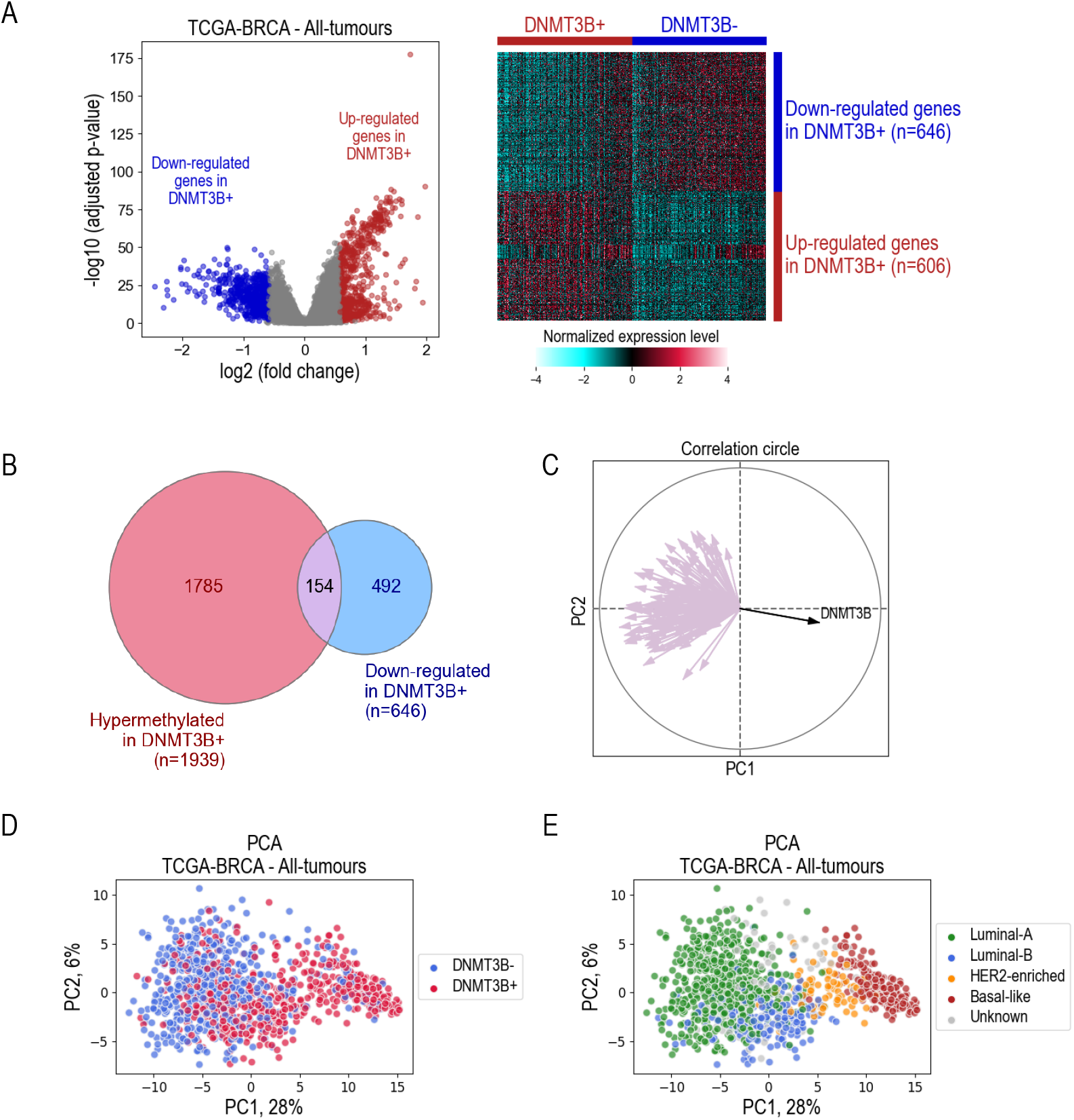
Genes regulated by aberrant *DNMT3B* activation. **A**: Results of the differential analysis of the expression profiles between the samples with high expression of *DNMT3B* (DNMT3B+) versus the samples with low expression of *DNMT3B* (DNMT3B-) in the TCGA-BRCA dataset, presented as a volcano plot and heatmap. Differentially expressed genes were identified using a t-test with an adjusted p-value threshold of < 0.05 and an absolute fold change > 1.5. In the volcano plot, the downregulated genes in DNMT3B+ versus DNMT3B-are shown as blue dots. The upregulated genes are shown as red dots. The heatmap shows hierarchical clustering, using Euclidean distance with Ward’s linkage for sample clustering and Pearson correlation for gene clustering. **B**: A Venn diagram illustrating the overlap between the results of differential analyses on methylation and expression data in the TCGCA-BRCA dataset. The red circle represents 1939 genes found to be significantly hypermethylated in DNMT3B+ tumours, while the blue circle contains 492 genes found to be significantly downregulated in these tumours. Of note, 154 genes were identified as belonging to both categories. **C**: The correlation circle illustrates the positions of 154 selected genes, together with *DNMT3B*, represented by arrows on the plot. These positions are mapped to the first two principal components from the PCA analysis performed on the whole transcriptome of the TCGA-BRCA dataset. **D**: Principal component analysis performed on the entire transcriptome of the TCGA-BRCA dataset. Blue dots correspond to DNMT3B-tumours. Red dots correspond to DNMT3B+ tumours. **E**: Same as **D** where the samples are coloured according to the corresponding molecular subtype of breast cancer.

Finally, for each of the 154 selected genes, we performed a univariate survival analysis using the Cox proportional hazards model and the log-rank statistical test to evaluate the impact of expression level of the gene on the risk of patients’ relapse in the eight breast cancer datasets. We considered the association with disease-free survival to be significant if both the Cox and the log-rank p-values were < 0.05. We found that for 28 out of 154 genes (18.2%), upregulation of expression levels was significantly associated with the probability of disease-free survival in at least three out of eight datasets: *ADCY1, AFF3, BCL2, C3orf18, CFB, CHAD, CST3, DNALI1, DUSP4, EGR3, FBP1, GASK1B, GATA3, GFRA1, GJA1, GREB1, IGFBP4, LRG1, NME5, PDE4A, PDZK1, PTPRT, SCNN1A, SCUBE2, SERPINA1, SRARP, TBC1D9, TFAP2B*. Figure 3A shows the hazard ratios (HR) obtained from Cox model for these genes in different datasets.

**Figure 3.**
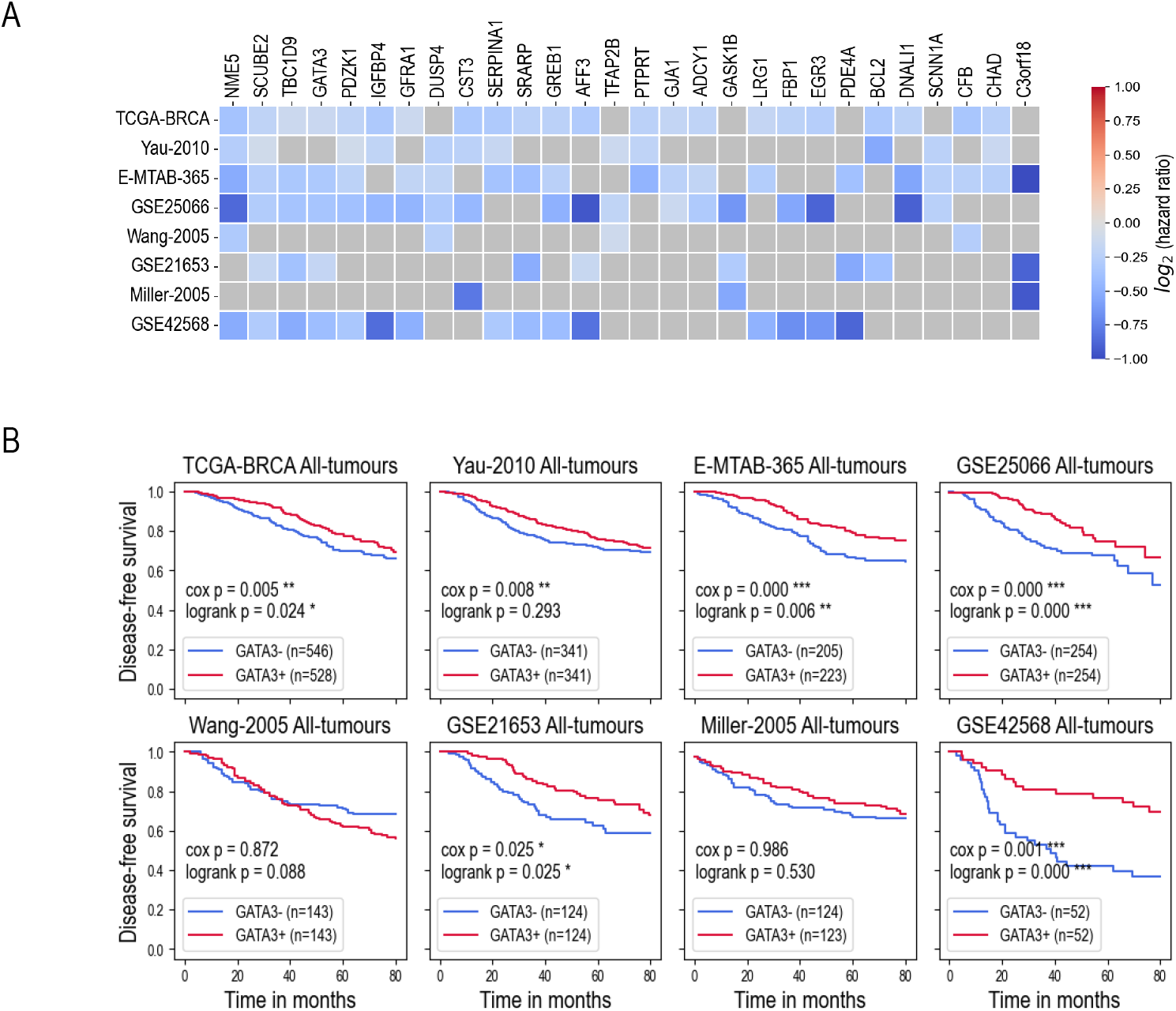
Aberrant expression of *DNMT3B* drives tumour suppressor gene inactivation. **A**: Results of the survival analysis performed on 154 selected genes in eight breast cancer datasets. The figure shows 28 genes for which the aberrant activation of *DNMT3B* was found to be significantly associated with patient prognosis in at least 3 out of 8 datasets. The x-axis corresponds to the genes, the y-axis shows the datasets. The value of log_2_(hazard ratio) calculated from the Cox proportional hazard model, for which the significant p-value < 0.05 was obtained, is shown in the blue colour map for each gene and dataset. The grey squares correspond to non-significant results. **B**: Kaplan-Meier survival curves across eight public datasets, calculated for two groups of patients based on *GATA3* expression levels: tumours with low *GATA3* expression (GATA3-, blue line) and tumours with high *GATA3* expression (GATA3+, red line). The threshold separating the groups corresponds to the median expression level. P-values are calculated using the log-rank test and the Cox proportional hazards model and are indicated by the following significance symbols: * for p < 0.05, ** for p < 0.01, and *** for p < 0.001.

For all 28 genes, the hazard ratios (HR) were found below 1.0, meaning that the overexpression of these genes predicts a lower risk of relapse for patients. These results suggest that aberrant activation of *DNMT3B* in breast cancer may specifically target hypermethylation of the promoter regions of genes responsible for anti-tumour activity.

Detailed results of survival analyses are shown in Figure 3B for the gene *GATA3* as an example. In addition, Supp. Figure S1 shows the results of a correlation analysis between the expression levels of *DNMT3B* and *GATA3* in eight publicly available datasets. In seven cases (out of 8), these correlation studies revealed a significant correlation between increased *DNMT3B* expression and a corresponding down-regulation of *GATA3* expression.

### The onco-suppressor gene *GATA3* is one of the main targets of epigenetic silencing controlled by *DNMT3B* in aggressive forms of breast cancer

Our results presented above suggest that the aberrant activation of *DNMT3B* in breast cancer induces a hypermethylation of CpG rich DNA regions in the promoters of several onco-suppressor genes, responsible for the more aggressive profiles of tumour cells. In fact, hypermethylation regulated by *DNMT3B* is one of the mechanisms used by tumour cells to silence onco-suppressor genes. However, we hypothesised that the same target genes could also be affected by mutations associated with a loss of function (LOF) or deleterious large genomic deletions (copy number alterations, CNA) leading to their aberrant inactivation. This idea is illustrated in Figure 4A. Therefore, we analysed copy number deletion and mutation data in the TCGA-BRCA dataset available on cBioPortal (https://www.cbioportal.org).

**Figure 4.**
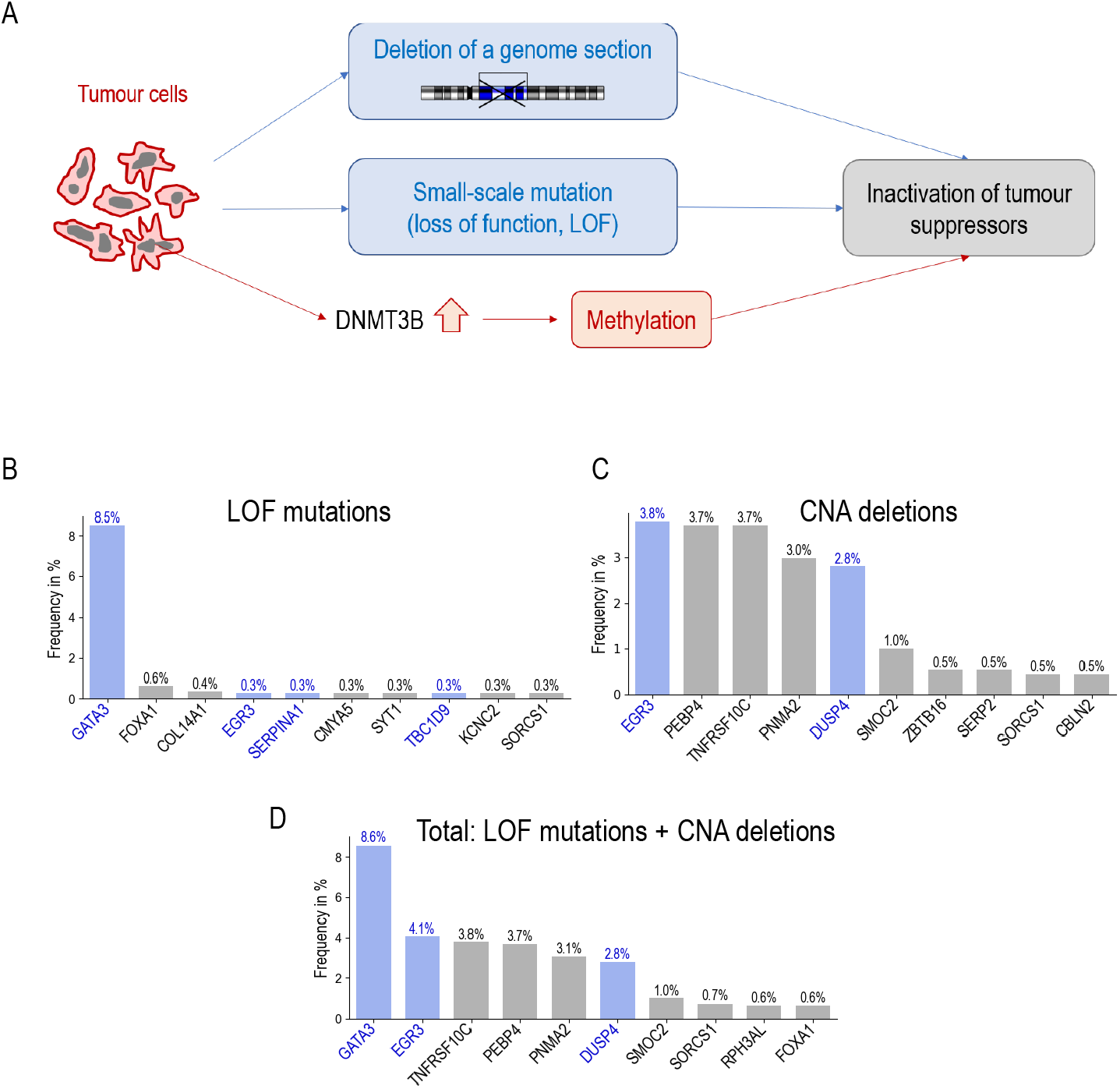
*GATA3* is a primary target for gene silencing in breast cancer. **A**: Possible mechanisms of tumour suppressor gene silencing in breast cancer. **B-D**: Frequency of loss of function (LOF) mutations (B), deleterious copy number alterations (C) and total frequency (D) calculated in the TCGA-BRCA dataset for 154 selected genes. The figure shows the top 10 genes ranked by frequency in descending order.

For the mutation data in the TCGA-BRCA dataset, we selected the most common types of gene mutations that result in a loss of function (LOF) of the corresponding protein: “stop gained”, “start lost”, “frameshift variant”, “in-frame deletion”, “in-frame insertion” and “protein altering variant”. Some other types of mutations, such as “missense variant” can also sometimes silence genes, but without detailed information on their functional consequences, we have not retained these mutations. Our aim was to identify, from our selection of 154 genes, those most frequently affected by LOF mutations in breast cancer. The calculated frequency of LOF mutations is shown in Figure 4B. We found that the gene *GATA3* was most frequently targeted by loss-of-function mutations in at least 8.5% of breast cancers. The other 153 genes were affected by LOF mutations in less than 1% of breast cancers. We repeated the same analysis for copy number alterations (CNA) with deletions of genomic regions (Figure 4C). The genes most frequently affected by CNA deletions in breast cancer were *EGR3* (3.8%), *PEBP4* (3.7%) and *TNFRSF10C* (3.7%). When the frequencies of LOF mutations and CNA deletions were added together (Figure 4D), the gene most frequently targeted for gene inactivation by these two mechanisms in breast cancer was *GATA3* (8.6%).

Finally, our discovery of epigenetic inactivation of the *GATA3* gene in addition to its genetic silencing makes the actual proportion of *GATA3* inactivation impressive: 8.6% of genetic silencing + 6.8% of epigenetic silencing = 15.4% (see details in the next sections). However, before confirming this hypothesis, we needed to verify that *GATA3* can be inactivated either genetically or epigenetically. These different modes of *GATA3* inactivation should therefore occur independently in different cells. To confirm this prediction, we have shown that these modes of gene inactivation occur independently and that there is only a small overlap between cells with each type of *GATA3* inactivation (see below).

### The downregulation of *GATA3* by aberrant activation of *DNMT3B* was confirmed in our independent cohort

To confirm the hypothesis of *GATA3* silencing by *DNMT3B*-mediated DNA methylation in its promoter region, we established an independent validation cohort of 19 breast cancer patients from the Centre Léon Bérard, Lyon, France (biobank CRB-CLB BB-0033-00050, dataset registered as CMT 2023-22_BS2023_-010_R201-004-415). For each frozen tissue sample, co-extraction of DNA and RNA was performed. We then generated bulk RNA-seq and methylation data on the Illumina MethylationEPIC v2.0 bead chip. We performed correlation analyses between average DNA methylation and expression levels of the genes *DNMT3B* and *GATA3* in the public TCGA-BRCA dataset and in our validation dataset of 19 patients. As shown in Figure 5, the expression level of *DNMT3B* is significantly correlated with the average level of DNA methylation in the promoter region of *GATA3*. This result was first demonstrated in the TCGA-BRCA dataset (Figure 5A) and then successfully confirmed in our validation cohort (Figure 5D). Furthermore, as expected, our results show that the expression level of *GATA3* is significantly anti-correlated with the average DNA methylation level in its promoter region in both datasets (Figure 5B and Figure 5E). Finally, a significant anti-correlation between the expression levels of *GATA3* and *DNMT3B*, initially observed in the TCGA-BRCA dataset (Figure 5C), was confirmed in our validation cohort (Figure 5F), suggesting that *GATA3* may indeed be down-regulated by the aberrant activation of *DNMT3B* in breast cancer.

**Figure 5.**
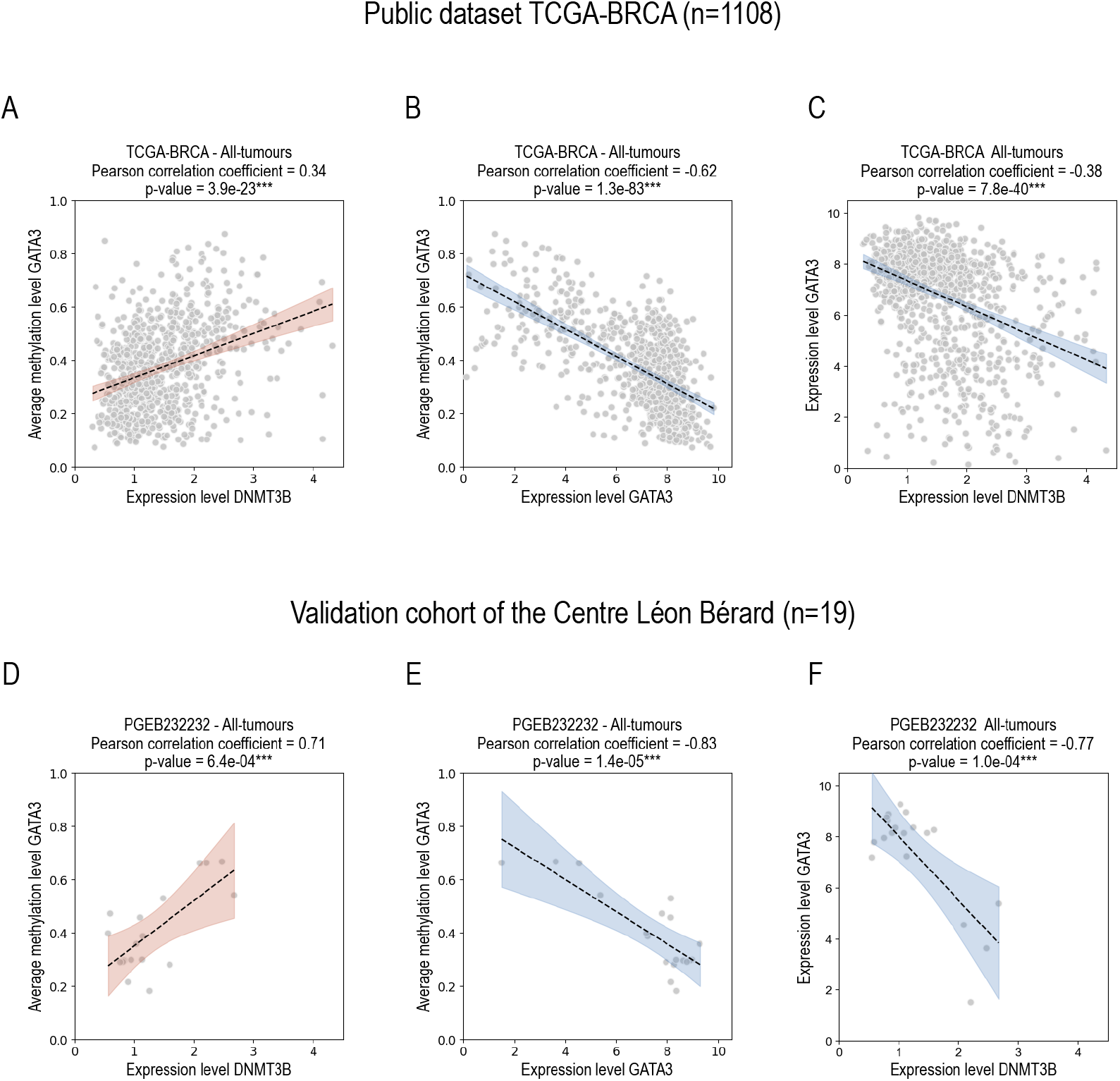
*GATA3* is regulated by *DNMT3B*-mediated DNA methylation in its promoter region. Results of correlation analyses performed in the TCGA-BRCA dataset and in our independent cohort of 19 breast cancer patients from the Centre Léon Bérard, Lyon, France. The average DNA methylation in the promoter region of *GATA3* is significantly correlated with the expression level of *DNMT3B* in both datasets (A and D). The expression level of *GATA3* is regulated by the methylation level in its promoter region (B and E). The expression levels of *GATA3* and *DNMT3B* are significantly correlated (C and F). In all plots we consider the correlation to be significant for p-value < 0.05, as indicated by the following significance symbols: * for p < 0.05, ** for p < 0.01, and *** for p < 0.001. The shaded area corresponds to the 95% confidence interval: in red if the p-value is significant and the correlation coefficient is positive, in blue if the p-value is significant and the correlation coefficient is negative, in grey if the p-value is not significant.

### GSEA analysis reveals common and distinct molecular signatures between DNMT3B+ tumours and tumours with *GATA3* mutations

Following the hypothesis that the tumours with ectopic activation of *DNMT3B* target the silencing of a number of onco-suppressor genes, primarily *GATA3*, we performed a Gene Set Enrichment Analysis (GSEA) in two different contexts, separately. On the one hand, we compared DNMT3B+ tumours inducing hypermethylation of the *GATA3* promoter region with DNMT3B-tumour samples and, on the other hand, tumours with genetic alteration of *GATA3* (LOF mutations or copy number deletions) with the same DNMT3B-tumour samples without LOF mutations or deleterious *GATA3* copy number alterations. The GSEA analysis developed by the Broad Institute (Subramanian et al., 2005; Mootha et al., 2003) determines whether an a priori defined set of genes shows statistically significant differences between two biological states. This method focuses on gene sets that share common biological functions. More than 10000 published gene sets are publicly available for GSEA analysis in several collections of the MSigDB database (Liberzon et al., 2011, 2015) on the Broad Institute website (https://www.gsea-msigdb.org/gsea/msigdb). For our analysis we selected the H (hallmarks), C2 (curated gene sets) and C5 (ontology gene sets) collections from the MSigDB database.

In the first GSEA analysis, we compared the DNMT3B+ tumours (with downregulation of *GATA3* expression and high levels of methylation in the promoter region of the gene - this group was named GATA3-METH) with the control group of tumour samples in which *DNMT3B* was not ectopically activated and no genetic alteration for *GATA3* was found (the control group was named GATA3-CTRL). The GATA3-METH group represents 6.8% of all breast cancer samples in the TCGA-BRCA dataset. In the second analysis, we compared the group of tumours with LOF mutations or copy number deletions of *GATA3* (this group was called GATA3-LOFDEL) with the same control group of tumours GATA3-CTRL. The GATA3-LOFDEL group represents 8.6% of all breast cancer samples in the TCGA-BRCA dataset. Figure 6A-B show the average methylation level of the *GATA3* gene in all three groups, a PCA projection, and the corresponding survival curves. The PCA projection shows that the GATA3-METH and GATA3-LOFDEL tumours correspond to two well separated populations. Disease-free survival was significantly shorter in the GATA3-METH and GATA3-LOFDEL groups than in the GATA3-CTRL group. Furthermore, we observed a strong correlation between the *GATA3* inactivation groups, i.e. GATA3-METH and GATA3-LOFDEL, and molecular subtypes (Figure 6B-C). In the GATA3-METH group, the majority of tumours belong to the basal-like molecular subtype (80.3%), 18.3% of tumours are HER2-enriched, only 1.4% of tumours are luminal-A, and no luminal-B tumours (Figure 6B). In contrast, the majority of tumours in the GATA3-LOFDEL group belong to the luminal molecular subtype (56.3% for luminal-A and 40.2% for luminal-B), while only 3.4% of tumours are HER2-enriched, and no basal-like tumours (Figure 6C). However, both groups with *GATA3* inactivation, whether by methylation, loss-of-function (LOF) mutations or deleterious CNA, are associated with a worse relapse prognosis compared to tumours in which *GATA3* is expressed and not mutated, regardless of the specific inactivation mechanism.

**Figure 6.**
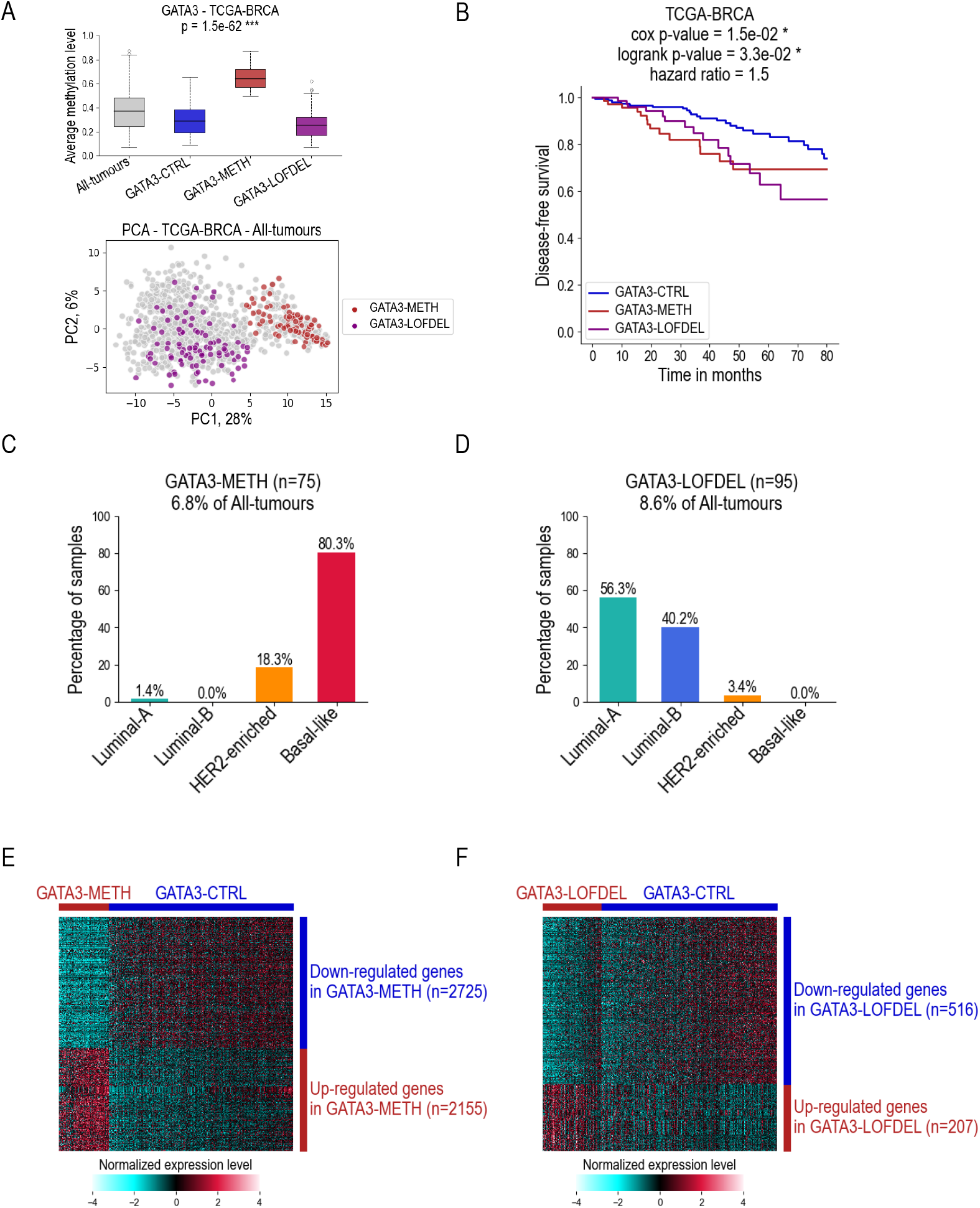
Results of differential analyses of transcriptomic data. **A**: Box plots of average methylation levels of the promoter region calculated for the gene *GATA3* in different groups of samples in the TCGA-BRCA dataset: whole dataset (All-tumours), control group (GATA3-CTRL), group in which *GATA3* is inactivated by methylation (GATA3-METH), group in which *GATA3* is inactivated by loss-of-function mutations or deleterious copy number alterations (GATA3-LOFDEL). The distribution of GATA3-METH and GATA3-LOFDEL samples is also shown in a PCA projection calculated with the same method as in Figure 2. **B**: Kaplan-Meier survival curves calculated for the GATA3-CTRL, GATA3-METH, and GATA3-LOFDEL tumour groups in the TCGA-BRCA dataset. Survival analysis is performed using the log-rank test and the Cox proportional hazards model. **C-D**: Percentage of tumour samples according to their molecular subtypes in the TCGA-BRCA dataset, for the GATA3-METH group, where *GATA3* is inactivated by methylation (C), and for the GATA3-LOFDEL group, where *GATA3* is inactivated by LOF mutation of a deleterious CNA (D). **E-F**: Results of differential analyses of total transcriptomic data in the TCGA-BRCA dataset between the GATA3-METH (C), GATA3-LOFDEL (D) and GATA3-CTRL tumour groups. Differentially expressed genes were identified using a t-test with an adjusted p-value threshold of < 0.05 and an absolute fold change > 1.5. The heatmaps show hierarchical clustering, using Euclidean-based distance with Ward’s linkage for sample clustering and Pearson correlation for gene clustering.

We then performed two differential expression analyses for GATA3-METH versus GATA3-CTRL and for GATA3-LOFDEL versus GATA3-CTRL (Figure 6E-F). In both experiments, for all genes of the genome, we calculated the corresponding fold changes of the average expression levels between the two compared groups. We then ordered all the genes by the obtained fold change values, from highest to lowest, and performed the subsequent GSEA analyses to identify significantly enriched or depleted gene sets for which the obtained nominal p-value < 0.05 and FDR < 0.25. The results of the GSEA analyses are shown in Figure 7 and Figure 8.

**Figure 7.**
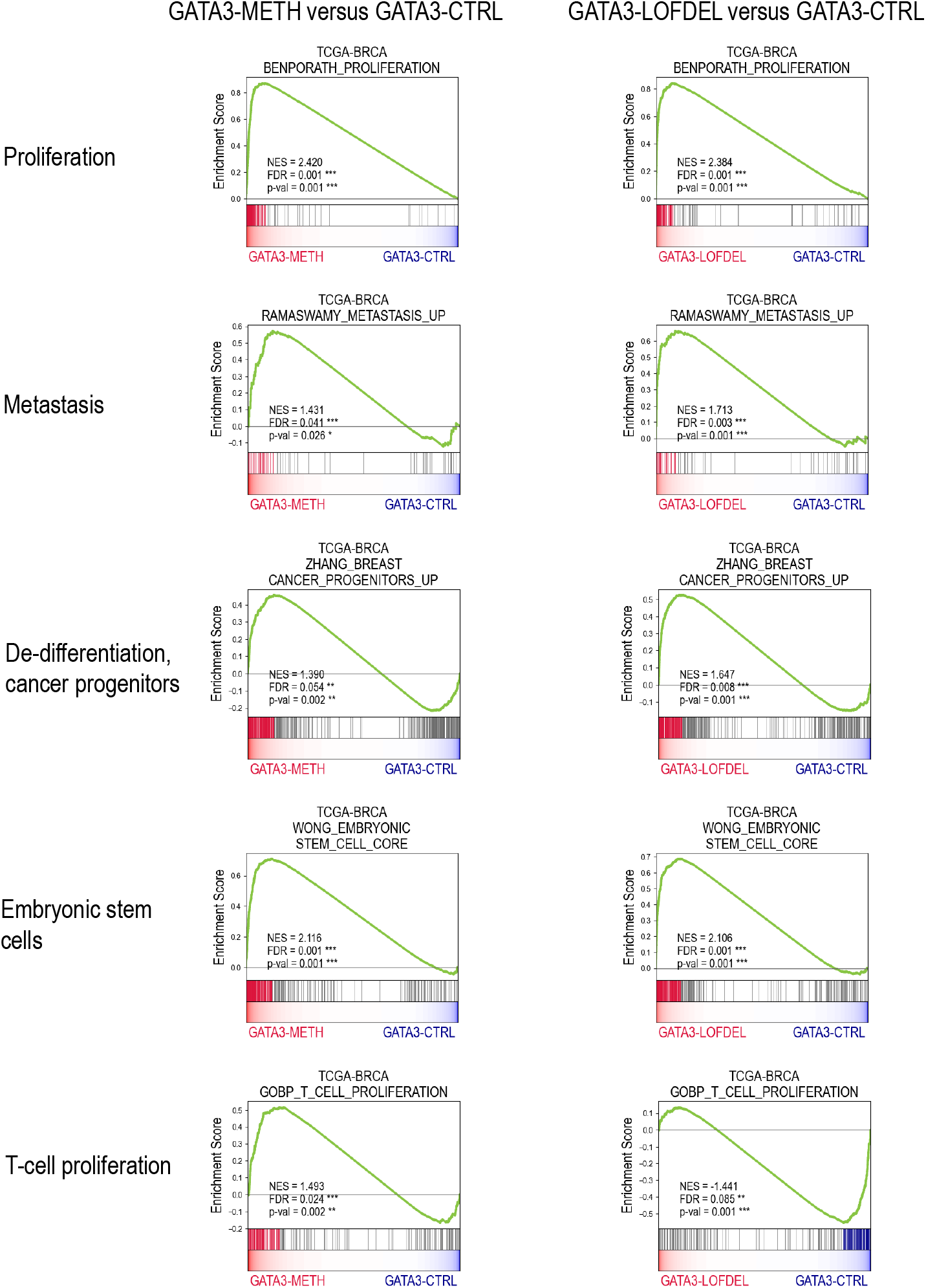
Main results of the GSEA analysis for transcriptomic profiles of GATA3-METH and GATA3-LOFDEL versus GATA3-CTRL tumours. Results of the GSEA analysis for transcriptomic profiles of GATA3-METH versus GATA3-CTRL, and GATA3-LOFDEL versus GATA3-CTRL tumours in the TCGA-BRCA dataset. GSEA plots illustrate the main enrichment/depletion profiles. For all the gene sets, the enrichment or depletion was considered significant with a nominal p-value < 0.05 and FDR < 0.25. The gene sets were selected from the Broad Institute’s MSigDB database of the (MSigDB collections C2, C5 or H).

**Figure 8.**
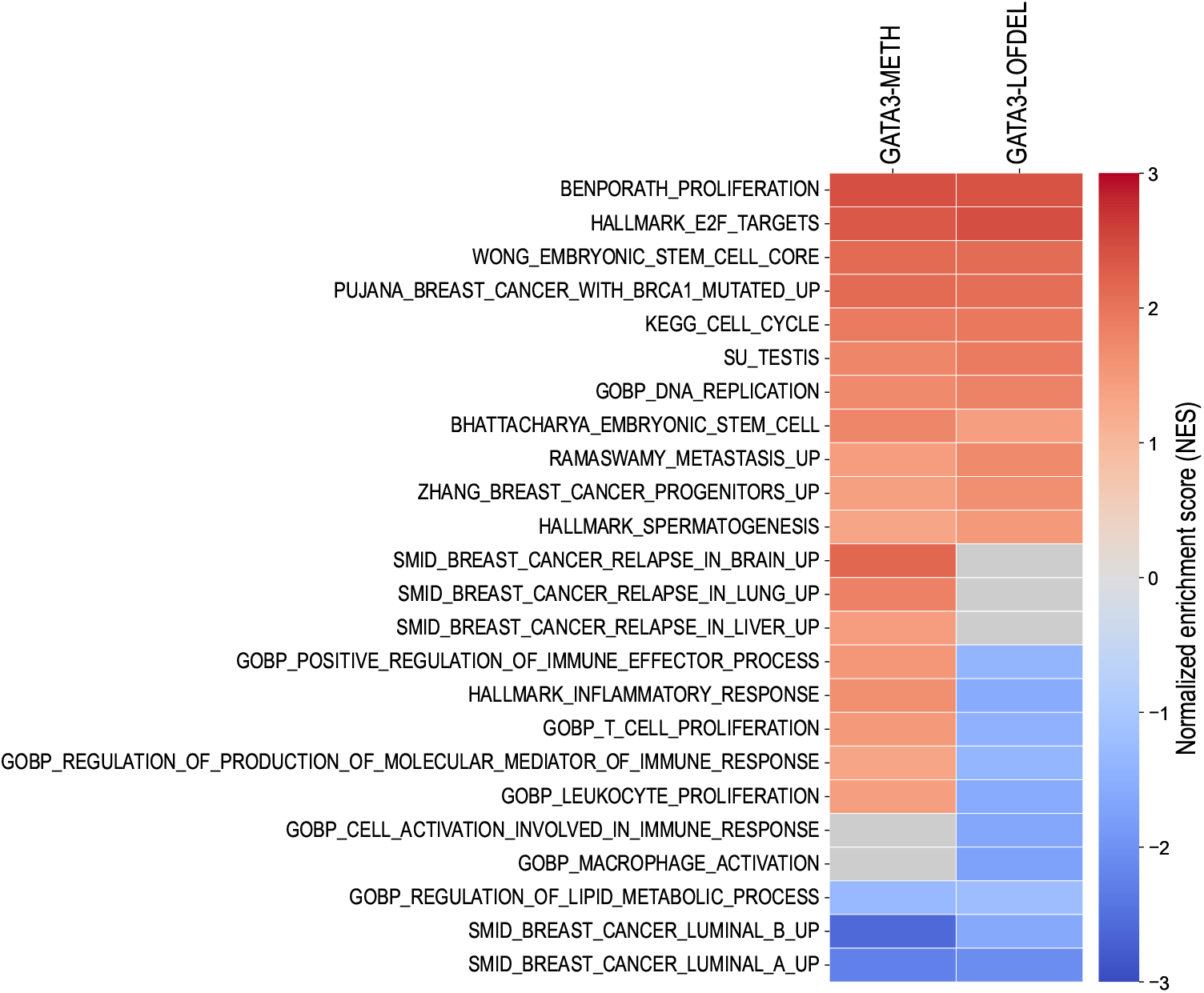
Gene Set Enrichment Analysis (GSEA) shows shared and distinct molecular signatures in the GATA3-METH and GATA3-LOFDEL tumours. The heatmap shows the normalised enrichment score (NES) obtained from the GSEA analysis of GATA3-METH vs. GATA3-CTRL, and GATA3-LOFDEL vs. GATA3-CTRL tumours in the TCGA-BRCA for different genes sets. Significantly enriched gene sets are shown in red, and significantly depleted gene sets are shown in blue. For all the gene sets, the enrichment or depletion was considered significant with a nominal p-value < 0.05 and FDR < 0.25. Grey cells correspond to non-significant results.

As expected, we found that the breast tumours in both groups, GATA3-METH and GATA3-LOFDEL, shared many molecular signatures associated with the aggressive properties of breast cancer. For example, both GATA3-METH and GATA3-LOFDEL tumours were significantly enriched in molecular signatures of proliferation, cell cycle acceleration, DNA replication, and metastatic progression. In addition, we found that these tumours significantly overexpressed the genes corresponding to the signature of breast cancer stem cell progenitors, suggesting that both *DNMT3B* activation and deleterious somatic alterations of *GATA3* promote a dedifferentiation of cancer cells (or proliferation of progenitor cells). Interestingly, both tumour subtypes GATA3-METH and GATA3-LOFDEL are significantly enriched in the molecular signature of mutated *BRCA1*. They are also both enriched in testis-specific and embryonic stem cell-specific gene expression signatures. The GATA3-METH and GATA3-LOFDEL cancer subtypes are also both significantly depleted in genes associated with luminal-A and luminal-B molecular subtypes, consistent with their aggressive profiles. The regulation of lipid metabolism is another common signature that is significantly depleted in both groups, indicating a disruption of lipid metabolism in these tumours.

Although many molecular signatures, especially those associated with aggressive forms of breast cancer and poor prognosis, are shared by both subtypes GATA3-METH and GATA3-LOFDEL tumours, there are also some differences. GATA3-METH tumours appear to be more prone to distant metastases in the brain, lung and liver than GATA3-LOFDEL tumours. One striking difference concerns the molecular signatures of inflammatory and immune responses. Several gene sets related to inflammatory response, T-cell and leukocyte proliferation, macrophage activation and immune response are significantly depleted in GATA3-LOFDEL tumours (Figure 8), whereas the majority of these gene sets are significantly enriched in the GATA3-METH group. These results suggest that GATA3-LOFDEL tumours are able to escape the immune response more efficiently than GATA3-METH tumours.

## Discussion

We investigated the role of ectopic activation of the DNA methyltransferase 3 beta (*DNMT3B*) in breast cancer. *DNMT3B* is normally expressed specifically in embryonic stem cells (ESC) and testis, but is absent or expressed at low levels in healthy breast tissues. This work was strongly motivated by our recent findings of aberrant activations of normally silent tissue-specific genes in breast cancer, in which *DNMT3B* was identified as one of the most potent prognostic biomarkers, along with four other ESC-specific genes, *EXO1, MCM10, CENPF* and *CENPE*, and robustly associated with a high risk of patients’ relapse (Jacquet et al., 2023). The gene *DNMT3B* was of particular interest because of its essential role in DNA *de novo* methylation. We hypothesised that the aberrant activation of *DNMT3B* could lead to aberrant DNA methylation in CpG-rich promoter regions of onco-suppressor genes, thereby silencing these genes by an epigenetic mechanism, in addition to other ways of targeting these genes directly by genetic alterations such as mutations or copy number changes.

Simultaneous analysis of RNA-seq transcriptomic and methylomic data from the TCGA-BRCA dataset reveals that the aberrant activation of *DNMT3B* in breast tumours induces a significant increase in promoter methylation for 1939 genes, which is associated with the down-regulation of expression levels for 646 genes. In the intersection of these gene lists, we identified 154 genes for which increased promoter methylation was associated with decreased expression levels as a result of *DNMT3B* activation. In addition, univariate survival analysis performed separately for each of the 154 genes shows that downregulation of the expression level of 28 out of 154 genes (18.2%) is significantly associated with a poor prognosis for disease-free survival. These results suggest that aberrant activation of the *DNMT3B* gene specifically targets the silencing of tumour suppressor genes.

In support of this conclusion, several of the 28 genes identified are known in the literature to be associated with anti-tumour activity, acting as onco-suppressors in cancer. For example, the gene *ADCY1*, found in our list of 28 genes, belongs to the adenylate cyclase (AC) pathway, which is related to the anti-tumour activity of xanthohumol (XN) against tumour cells, regulating various cellular functions via the activation of protein kinase A (PKA)-dependent phosphorylation (Shikata et al., 2017). Other examples include the genes *EGR3, GATA3* and *NME5*, which were also independently identified in our analysis. Overexpression of the *EGR3* gene is known to inhibit the growth of hepatocellular carcinoma (Zhang et al., 2017; Wang et al., 2017), and is also associated with a better prognosis in patients with gastric cancer (Liao et al., 2013) and prostate cancer (Pio et al., 2013). The gene *GATA3* is a transcription factor involved in the differentiation of mammary epithelial cells and is able to suppress epithelial-to-mesenchymal transition. Low levels of *GATA3* expression have been associated with tumour progression and poor patient prognosis in breast cancer (Reiswich et al., 2023). The gene *NME5* is a member of the nonmetastatic (NME) gene family and is predicted to be involved in the negative regulation of the oxidative stress-induced intrinsic apoptotic signalling pathway and spermatid development. Increased expression of *NME5* has been associated with a prolonged survival in breast cancer and may play a role as a tumour suppressor gene (Wu et al., 2020).

Methylation of gene promoter regions is one of several possible mechanisms by which tumour cells can silence a gene. However, the tumour can also achieve a similar goal by making specific mutations in the DNA sequences of a selected number of genes, leading to a loss of function (LOF) of these genes. In addition, deletions of large regions of the genome could also lead to the corresponding gene copy number alterations (CNA). To determine which genes are the most common targets for LOF mutations and deleterious CNA in breast cancer, we calculated the frequencies of these alterations for 154 genes in the TCGA-BRCA dataset. We found that the most frequent DNA alterations among the 154 genes occur in the gene *GATA3*, which is mutated with a LOF in at least 8.5% of the tumour samples. Large genomic deletions are not common for this gene, less than 0.1%. Overall, the gene *GATA3* is altered in 8.6% of tumour samples when both LOF mutations and CNA deletions are taken into account. The second gene frequently altered in breast cancer is *EGR3*. This gene is the target of CNA deletions in 3.8% of tumour samples and only 0.3% of LOF mutations, for a total of 4.1% of alterations. The other genes in our selection are altered with LOF mutations or CNA deletions in less than 4% of tumours.

According to our results, the gene *GATA3* appears to be the main target for silencing in the DNMT3B+ aggressive forms of breast cancer, with twice the frequency of deleterious somatic alterations compared to the other genes in our selection. *GATA3* is known to be the most highly expressed transcription factor in the mammary epithelium, and is involved in growth control and maintenance of the differentiated state in luminal epithelial cells of the adult mammary gland (Usary et al., 2004; Kouros-Mehr et al., 2006, 2008). In breast cancer, *GATA3* drives invasive tumour cells to undergo epithelial-mesenchymal transition reversal, leading to suppression of cancer metastasis (Yan et al., 2010). Several studies have reported frequent somatic mutations of *GATA3* up to 18% in breast cancer, predominantly in the ER-positive molecular subtypes (Banerji et al., 2012; Ellis et al., 2012; Cancer Genome Atlas Network, 2012; Bianco et al., 2022). However, both wild-type and mutant products of *GATA3* can be expressed and, depending on the nature of the mutation, the effects of *GATA3* can shift from tumour suppressing to tumour promoting activities during tumorigenesis (Cohen et al., 2014; Takaku et al., 2015). In contrast, silencing by DNA methylation of the *GATA3* promoter only leads to increased tumour aggressiveness due to loss of *GATA3* expression. Accordingly, loss of *GATA3* expression has been reported to be associated with basal-like breast cancer aggressiveness, *HER2* overexpression, oestrogen and progesterone receptor negativity, and reduced survival (Granit et al., 2013; Tkocz et al., 2012; Chu et al., 2012; Reiswich et al., 2023).

Our GSEA analyses showed that the DNMT3B+ tumours with hypermethylation of the promoter region of *GATA3* (GATA3-METH group) and the tumours with deleterious somatic alterations of *GATA3* (GATA-LOFDEL group) shared the same molecular signatures related to breast cancer aggressiveness: proliferation, cell cycle, DNA replication, metastasis, aberrant activation of tissue-specific genes and enrichment in breast cancer precursors. Interestingly, both GATA3-METH and GATA-LOFDEL groups of tumours were significantly enriched in the gene signature of mutated *BRCA1*. Interestingly, Tkocz et al. (2012) showed that *GATA3* interacts with a C-terminal region of *BRCA1* and that this interaction is important for normal breast differentiation as well as for the repression of genes associated with triple-negative and basal-like breast cancer. In addition, Bai et al. (2021) showed that *BRCA1* depletion stimulates methylation of the *GATA3* promoter, thereby repressing *GATA3* transcription, and that restoration of *GATA3* expression in *BRCA1*-deficient tumour cells activates mesenchymal-epithelial transition, suppressing tumour initiation and metastasis.

We have shown that the downregulation of *GATA3* in breast cancer can be controlled by an aberrant activation of *DNMT3B*, which induces aberrant methylation in the promoter region of *GATA3*, in the absence of deleterious somatic alterations. This mechanism allows breast cancer cells with particularly aggressive characteristics to significantly increase the risk of patient relapse. Our findings suggest that this group of breast cancers could potentially be targeted with a novel therapeutic approach aimed at reactivating *GATA3* through DNA methylation reprogramming or the use of DNA demethylating agents.

## Conclusions

We discovered that aberrant activation of *DNMT3B* in breast cancer regulates the methylation of the promoter region of *GATA3* and can repress this gene. When *DNMT3B* is activated, it acts similarly to a somatic loss-of-function mutation on *GATA3* or a copy number deletion. As a result, *GATA3* is downregulated in tumour cells. In aggressive breast tumours where *GATA3* is not expressed and not mutated, a new targeted therapy based on demethylation of the promoter region of *GATA3* could be considered.

## Methods

### Publicly available normal tissue and breast cancer datasets

To obtain expression profiles in normal tissues, we used RNA sequencing (RNA-seq) data provided by the GTEX portal and the NCBI Sequence Read Archive (datasets PRJNA280600, PRJEB4337, PRJEB2445, PRJNA270632, GSE70741, GSE53096). We also used 8 breast cancer datasets with available survival data from public data repositories: GDC Data Portal, Array-Express, NCBI GEO and USCS Xena. A detailed description of the datasets can be found in Supp. Table S3.

The RNA-seq data of normal human tissues from the GTEX repository and the NCBI Sequence Read Archive contain 2955 samples from 44 different tissues: 2913 samples from 39 adult tissues and 5 samples from embryonic stem cells. Some tissues were pooled in more general tissue groups. In total, we obtained 26 tissue groups: 18 tissue groups for adult tissues and 1 tissue group for embryonic stem cells.

The transcriptomic data of the microarray datasets E-MTAB-365, GSE25066, GSE21653, GSE42568, Miller-2005, Wang-2005 and Yau-2010 were obtained with Affymetrix Human Genome Arrays U133 Plus 2.0, U133A and U133B. The data were normalised using Robust Multi-array Average (RMA) method and then log-transformed. For the TCGA-BRCA dataset, we used the RNA-seq values normalised by FPKM method provided directly from the GDC Data Portal. The FPKM values were log-transformed by taking log_2_(1+FPKM). For normal tissue RNA-seq datasets, we downloaded pre-processed raw counts and normalised them to logtransformed RPKM units.

For the TCGA-BRCA dataset, we also downloaded methylation data (beta values from the Illumina Human Methylation 450 bead chip) for 766 samples from the GDC portal. The copy number alteration and mutation data from the TCGA-BRCA dataset were downloaded from cBioPortal (https://www.cbioportal.org).

### Validation cohort of 19 breast cancer patients

Breast cancer samples from 19 patients were provided by the biobank of the Centre Léon Bérard, Lyon, France (CRB-CLB BB-0033-00050). The data set is registered as CMT 2023-22_BS2023_010_R201-004-415. All patients were informed and did not object to their data and samples being used for this study. Tumours were snap frozen in liquid nitrogen at the time of surgical resection, after pathological review, and stored at the respective hospital’s biological resource centre. Corresponding pathological, clinical, and follow-up data were obtained from the collection centre.

#### Sample preparation

Frozen tissue samples were used for the simultaneous extraction of DNA and RNA using the AllPrep DNA/RNA Kit (Qiagen®). The extracted nucleic acids were then processed to generate bulk RNA-Seq and DNA methylation data.

#### RNA-Seq data

RNA sequencing libraries were prepared using the NEBNext® Ultra™ II Total RNA Library Prep Kit (New England Biolabs, Ipswich, MA, USA) according to the manufacturer’s instructions. This protocol included a ribodepletion step to remove ribosomal RNA, thereby enhancing the detection of both coding and non-coding transcripts. Unique Dual Index (UDI) barcodes were used to multiplex samples during sequencing. Libraries were sequenced on the Illumina NovaSeqX+ platform in paired-end mode, generating 2 × 150 bp reads. A target of 35 million paired-end reads per sample (70 million total reads) was achieved to ensure sufficient coverage for downstream analyses. Reads were aligned to the UCSC hg38 genome using STAR (2.7.11b) (Dobin et al., 2013) and then counted using the HTSeq framework (2.0.5) (Anders et al., 2015). Read counts were calculated as reads per kilobase million (RPKM) and log transformed to log2(1+RPKM).

#### DNA methylation data

Bisulfite-treated DNA was hybridised to the Infinium MethylationEPIC™ v2.0 arrays (Illumina, San Diego, CA, USA). Raw DNA methylation intensity data files (IDAT) were processed with RnBeads 2.0 using default parameters (Müller et al., 2019) to generate matrices of beta values.

### Differential Analysis of Methylation Data

In the TCGA-BRCA dataset, the association between DNMT3B+ versus DNMT3B-groups and DNA methylation levels at each CpG site (EWAS) was calculated using the standard linear model (lm) function of R (3.6.1). P-values obtained for each probe along the genome were aggregated using the Comb-p v2 software (Pedersen et al., 2012) with the following options: -c 5 –seed 1e-5 –dist 1000 –region-filter-p 0.05 –region-filter-n 2.

### Statistical Analyses of Transcriptomic Data

In correlation analyses, to measure the linear relationship between two variables, we calculated the Pearson correlation coefficient and corresponding p-value with the two-sided alternative using the Python package “scipy” version 1.14.1. The linear regression fit and confidence interval were calculated using the Python package “statsmodels” version 0.14.4.

For survival analyses, the time from enrolment to relapse was considered as the disease-free survival endpoint. Patients without relapse were censored at the last follow-up. Survival analyses were performed using the log-rank statistical test to compare the survival probabilities between different patient groups, and the Cox proportional hazards model to assess the effect of a numerical variable on the risk of relapse. Survival estimates were obtained using the Kaplan–Meier method. Survival analyses were performed using the “lifelines” package, version 0.30.0 of Python.

Expression levels between different patient groups were compared using the ANOVA statistical test. The p-values obtained were adjusted for multiple comparisons using the Benjamini-Hochberg procedure. In all statistical analyses described in this section, a p-value < 0.05 was considered statistically significant.

### Gene Set Enrichment Analysis (GSEA)

The GSEA analysis was performed on the C2, C5 and H collections of gene sets provided by the Broad Institute in the MSigDB database (https://www.gsea-msigdb.org/gsea), using the GSEA software “gseapy” available on the website.

### Manuscript Preprint

A preprint version of this manuscript was created using the following Latex template: https://github.com/quantixed/manuscript-templates.

## List of abbreviations

CNA: Copy Number Alterations
CTA: Cancer Testis Antigens
ER: Oestrogen Receptor
ESC: Embryonic Stem Cell
EWAS: Epigenome-Wide Association Studies
FPKM: Fragments Per Kilobase of transcript per Million mapped reads
GDC: Genomic Data Commons Data Portal
GEC: Gene Expression Classifier
GEO: Gene Expression Omnibus
GTEX: Genotype-Tissue Expression (GTEx)
HER2: Human Epidermal Growth Factor Receptor 2
LOF: Loss of Function
NCBI: National Center for Biotechnology Information
PR: Progesterone Receptor
RMA: Robust Multi-array Average
RNA-Seq: RNA Sequencing
RPKM: Reads Per Kilobase Million
TCGA: The Cancer Genome Atlas
TNBC: Triple Negative Breast Cancer
USCS: University of California Santa Cruz

## Declarations

## Acknowledgements

We thank the Centre Léon Bérard (CRB-CLB BB-0033-00050) biobank in Lyon, for sharing human biological samples and Melinda Teyssier form Institutional Data Base unit for collecting clinical data.

## Funding

This research was supported by the Cancer ITMO [Multi-Organisation Thematic Institute of the French Alliance for Life Sciences and Health (AVIESAN)] MIC program. This work also received support from the “Association Espoir Isere contre le cancer” and from “Groupement des Entreprises Françaises dans la Lutte contre le Cancer (Gefluc)” attributed to EJ. Additional funding was provided by Plan Cancer Pitcher, MSD Avenir ERICAN programs as well as by the ANR EpiSperm4 and 5 and the INCa - IreSP programs. Data processing were performed using the CIMENT/GRICAD infrastructure (https://gricad.univ-grenoble-alpes.fr), which is supported by Grenoble research communities.

## Availability of data and materials

The majority of datasets analysed during the current study are publicly available. The corresponding identifiers are given in Supp. Table S3. The full data outputs supporting the conclusions of this article are included within the article and its additional files.

## Ethics approval and consent to participate

The material used in the study was collected in agreement with all applicable laws, rules, and requests of French and European government authorities, including the patient’s informed consent.

Samples were collected in the context of patient diagnosis and remainders conserved in the Biological Resource Center (BRC) of the Centre Léon Bérard (no. CRB-CLB BB-0033-00050) registered at the Ministry of Research (DC-2008-99 and AC-2019-3426). All patients were informed and did not object to their data and samples being used for this study. The BRC is quality certified ISO 20387 and ISO 9001 ensuring scientific rigour for sample conservation, traceability, and quality, as well as ethical rules observance, and defined rules for transferring samples for research. Sample usage is reviewed by the ethical review board of the Centre Léon Bérard before any transfer (approval n°for this study N° CMT 2024-22).

The samples are properly coded to ensure anonymity of the donors or usage of any clinical information and the data management was approved by the data protection officer of the CLB as conformed to the GDPR rules (n° R201-004-415).

## Competing interests

The authors declare that they have no competing interests.

## Consent for publication

Not applicable.

## Authors’ Contributions

SK, SR, and EBF designed the work. FC and EBF retrieved and processed raw methylation and transcriptomic data. ALV conducted the validation phase of the project. IJ performed preliminary bioinformatic analyses. SR and EJ performed the mining of the literature. EBF realized and interpreted the *in-silico* data analyses. STE prepared biological samples from the Centre Léon Bérard biobank. EBF wrote the first draft of the manuscript including the figures and output tables. SR and SK substantially revised the manuscript. All authors read and approved the final manuscript.

## Notes

### Competing Interest Statement

The authors have declared no competing interest.

