## Supplementary Figure S1 for "Discovery of Epigenetically Silenced Tumour Suppressor Genes in Aggressive Breast Cancer Through a Computational Approach"

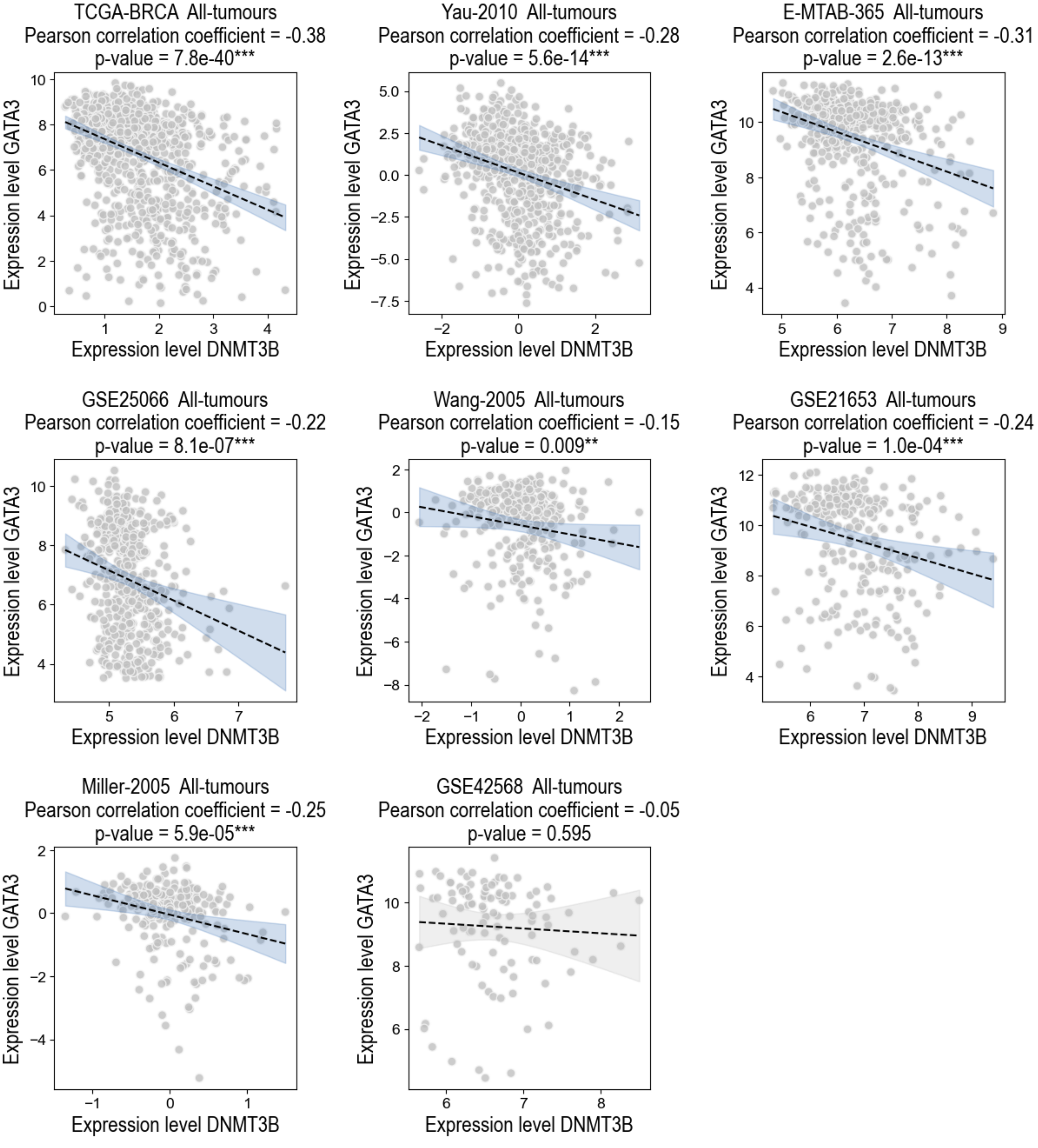


**Supplementary Figure S1.** Correlation plots between the expression levels of the genes DNMT3B and GATA3 in 8 public breast cancer datasets. The black dashed line shows the fitted regression. The shaded area corresponds to the 95% confidence interval: in blue if the corresponding p-value < 0.05, otherwise in grey.
